# Identification of an early hippocampal recognition system using intracerebral evoked potentials in humans

**DOI:** 10.1101/2022.11.02.513525

**Authors:** Víctor J. López-Madrona, Agnès Trébuchon, Ioana Mindruta, Emmanuel J. Barbeau, Andrei Barborica, Costi Pistol, Irina Oane, F. Xavier Alario, Christian G. Bénar

**Affiliations:** Aix Marseille Univ, INSERM, INS, Inst Neurosci Syst, Marseille, 13005, France; APHM, Timone Hospital, Epileptology and cerebral rhythmology, Marseille, 13005, France; APHM, Timone Hospital, Functional and stereotactic neurosurgery, Marseille 13005, France; Physics Department, University of Bucharest, 405 Atomistilor Street, Bucharest, Romania; Centre de Recherche Cerveau et Cognition, Université de Toulouse, Université Paul Sabatier Toulouse, Toulouse 31052, France; Centre National de la Recherche Scientifique, CerCo (UMR5549), Toulouse 31052, France; Aix-Marseille Université, CNRS, LPC, Marseille, France

## Abstract

The role of the hippocampal formation in memory recognition has been well studied in animals, with different pathways and structures linked to specific memory processes. In contrast, the hippocampus is commonly analyzed as a unique responsive area in most electrophysiological studies in humans, and the specific activity of its subfields remains unexplored. We combined intracerebral electroencephalogram recordings from epileptic patients with independent component analysis (ICA) during a memory recognition task involving the recognition of old and new images to disentangle the activities of multiple neuronal sources within the hippocampus. We identified two sources with different responses emerging from the hippocampus: a fast one (maximum at ∼250 ms) that could not be directly identified from raw recordings, and a later one, peaking at ∼400 ms. The earliest component was found in 12 out of 15 electrodes, with different amplitudes for old and new items in half of the electrodes. The latter component, identified in 13 out of 15 electrodes, had different delays for each condition, with a faster activation (∼290 ms after stimulus onset) for recognized items. We hypothesize that both sources represent two steps of hippocampal memory recognition, the faster reflecting the input from other structures and the latter the hippocampal internal processing. Recognized images evoking early activations would facilitate neural computation in the hippocampus, accelerating memory retrieval of complementary information. Overall, our results suggest that hippocampal activity is composed by several sources, including an early system for memory recognition, that can be disentangled with ICA methods.

**SIGNIFICANCE STATEMENT:** In the human memory circuit, the hippocampus is considered as a relatively late structure, associated to the retrieval of elaborated memories. In most electrophysiological studies, it is analyzed as a unique responsive area, and the specific activity of its subfields remains unexplored. In this work, we combined intracerebral recordings with independent component analysis to separate the electrophysiological activity from two different substructures of the hippocampus. We analyzed the responses of both sources in a memory task involving the recognition of old and new images. Our results revealed new hippocampal dynamics associated to different subfields, with memory recognition occurring much faster than previously reported. Importantly, we confirmed the potential of independent component analysis, which can be extended to other brain areas.

## INTRODUCTION

The interaction between the different regions of the hippocampal formation plays a key role in many memory processes (Bird and Burgess, 2008; Carr et al., 2011; Hainmueller and Bartos, 2020). Inputs from CA3 and the entorhinal cortex (EC) integrate distinct information to CA1. Previous studies on rats have related the trisynaptic pathway and the autoassociation network of CA3 with memory retrieval (Rolls, 2010), and EC with the process of environmental information and the encoding of memories (Hasselmo et al., 2002). In the dentate gyrus, inputs from two different streams of the EC (lateral and medial EC) converge, reflecting the “what” and the “where” pathways, respectively (Fernández-Ruiz et al., 2021). Despite the large knowledge of these structures in animals, their specific electrophysiological activities in humans remain comparatively much less well understood.

Intracerebral recordings (stereotaxic electroencephalography, SEEG) are considered an excellent approach to analyze the dynamics of the hippocampus, with a millisecond time scale and fine spatial specificity (Fernández et al., 1999; Halgren et al., 1995). Event-related potentials (ERPs) are elicited in the hippocampus during recognition of known faces or words, with larger amplitudes between 400 and 600 ms post-stimulus (hippocampal P600) for successful item recognition (Barbeau et al., 2008; Fernández et al., 1999; Trautner et al., 2004). The timing of this response, together with the activation of other brain areas, allows a rough estimation of the memory circuit of the brain. Early potentials are generated along the whole ventral pathway (Allison et al., 1999). However, the first structure showing a modulation by the old/new status of the stimuli is the perirhinal cortex, with higher responses for known items at ∼200-300 ms, followed by the supplementary motor area, and by various frontal and parietal regions (Despouy et al., 2020; Gonzalez et al., 2015). In this context, the hippocampus has been considered as one of the latest structures involved in recognition (Despouy et al., 2020), with the first difference between new and old elements occurring almost 500 ms after stimulus onset. This result has suggested that the hippocampus is not involved in the earliest stages of memory recognition, but rather in a slower process that retrieves complementary information (Despouy et al., 2020).

Despite the high spatial resolution of SEEG, it is extremely difficult to identify and analyze the different substructures of the hippocampus, namely CA1, CA3 or the DG. These structures and SEEG contacts are roughly of the same size — a few millimeters. A single sensor may spatially cover more than one structure, while others may not be sampled. In addition, SEEG sampling is constrained by clinical indications. Only a limited number of contacts are placed in the hippocampus and their location varies across patients. The different substructures of the hippocampus are folded one in the other, resulting in overlapped field potentials. Therefore, the hippocampus is most commonly analyzed as a single responsive area in SEEG, blurring the contribution of each substructure to the recordings (Barbeau et al., 2017; Ludowig et al., 2010).

When the activities of various structures overlap in time and space, independent component analysis (ICA) may be used to separate the time-courses of the different current generators contributing to the recorded field potentials (Herreras et al., 2022, 2015; Makarova et al., 2011). The efficiency of ICA to disentangle hippocampal pathways has been well established in animal studies, where it has helped isolating the different inputs to CA1 (Benito et al., 2016; López-Madrona et al., 2020) and to the DG (Benito et al., 2014; Fernández-Ruiz et al., 2021, 2013). In humans, methods such as bipolar montages or current source density analysis are commonly used to measure the local inflow and outflow of currents in a specific location (Mitzdorf, 1985; Nicholson and Freeman, 1975). However, these methods may not recover the correct time-courses of the local sources (Fernández-Ruiz and Herreras, 2013; Martín-Vázquez et al., 2013; Michelmann et al., 2018). For example, in the case of two co-localized sources (i.e., close to the same SEEG contact), the bipolar montage would measure a combination of the activities of both structures. This overlap can have critical consequences. If the sources have anticorrelated activities, one or both signals would be cancelled. ICA has been recently proposed as an optimal solution for re-referencing intracerebral EEG data and identifying neural generators in LFP recordings (Michelmann et al., 2018), an approach that has so far been restricted to animal studies.

Using such approach, we clearly identified for the first-time different sources of the electrophysiological signal in the hippocampus during a memory recognition task. First, we applied ICA on SEEG recordings, identifying two different hippocampal current generators and disentangling their time courses from other close or distant activities. Then, we examined the spatial topography of each hippocampal component and corroborated the local origin of the sources. Finally, we characterized the dynamics of the hippocampal responses during a memory recognition task.

## METHODS

### Participants

Ten patients (three females) with pharmacoresistant epilepsy volunteered for this study. They were implanted with SEEG depth electrodes along the longitudinal axis of the medial temporal lobe during presurgical evaluation. All these patients had been implanted in the hippocampus, either anterior or posterior, for clinical reasons. Five patients were recruited at La Timone hospital (Marseille, France) and the other five at the Emergency University Hospital Bucharest (Romania). None of the patients presented sclerosis. For those patients recorded in Marseille, the hippocampus exhibited the stereotyped responses during an oddball task, which confirmed the correct functionality of this structure. Table 1 shows the clinical information for each patient. This research has been approved by the relevant Institutional Review Board (Comité de Protection des Personnes, Sud-Méditerranée I, ID-RCB 2012-A00644–39) and the Bucharest University ethical committee approval (CEC 23/20.04.2019). All participants signed a written informed consent form regarding this research.

**Table 1:** Clinical information of each patient.

| ***Patient*** | **Age** | **Epilepsy** | **Language organization** | **Electrode/s location** |
| --- | --- | --- | --- | --- |
| 1 | 36 | Bilateral temporo-mesial | Atypical bilateral | Right anterior<br>Right posterior |
| 2 | 37 | Bilateral temporo-mesial | Left typical | Right anterior<br>Left anterior |
| 3 | 17 | Left Operculo-Insular | Left typical | Left anterior |
| 4 | 36 | Right temporo-mesial | Left typical | Right anterior<br>Left anterior |
| 5 | 26 | Bilateral extensive on heterotopia | Right atypical | Left anterior<br>Right posterior<br>Left posterior |
| 6 | 47 | Left temporo-mesial | Left typical | Left posterior |
| 7 | 38 | Right insular | Left typical | Right anterior |
| 8 | 27 | Left insular | Left typical | Left posterior |
| 9 | 23 | Right parieto-temporal<br>heterotopia | Left typical | Right anterior |
| 10 | 29 | Right temporo-occipital<br>heterotopia | Left typical | Right posterior |

### Experimental paradigm

Each block of the recognition memory task started with an encoding phase, during which 12 pictures were presented, one after the other, and the participant was asked to memorize them. Each picture was a simple colored drawing of a familiar item (e.g. a dog or a car) on a grey background. The picture database and precise selection criteria are described below. After a distracting video of one minute (silent excerpts from a documentary showing birds and landscapes), the recognition phase involved a set of 24 pictures from the same database. Half of these pictures had been presented during the encoding phase, while the other 12 were new never-presented pictures. Participants were asked to press two different buttons if they recognized the image as having been presented earlier during encoding (“old” condition), or if the image was “new” to them. The latency of this response is the reaction time (RT). Stimuli presentation and response logging were controlled by the software E-prime 3.0 (Psychology Software Tools, Pittsburgh, PA).

Each trial started with a fixation cross presented in the center of the screen for 1000 ms, followed by the experimental picture, presented for 1000 ms in the encoding sub-block and for 1500 ms in the recognition sub-block. The subsequent inter-trial interval was fixed to 1000 ms in both blocks. For each participant, a total of 7 blocks were programmed to be displayed consecutively, using different images.

We selected 24 × 7 = 168 images to be used as experimental materials from the database of Duñabeitia et al., 2018. They were selected as having high name agreement (above 90%). To ensure that the observations were not driven by item-specific properties, different experimental lists were created for each participant. The items were separated into two matched groups of 84 items to serve as old and new, alternatively across patients. Across the “old” and “new” groups of items there were roughly equal numbers of natural and artefact stimuli, with matched visual complexities; their names in French were matched for name agreement, length in syllables, and (log) lexical frequency of use (normative data from Duñabeitia et al., 2018 or New et al., 2004). The 84-item groups were further broken down into 7 groups to be used in the different blocks, with items matched for visual complexity and (log) lexical frequency across the 7 groups. All matching across picture groups was performed with the MATCH utility (van Casteren and Davis, 2006). In the encoding phases, the 12 items were presented in a random order; in the recognition phases, the items were presented in a pseudo-random order, with the constraint that there were never more than 3 “old” or “new” items in a row.

### SEEG recordings

SEEG recordings were performed using depth electrodes, implanted in stereotactic conditions (Talairach et al., 1992; Alcis, Besançon, France, and Dixi Medical, Chaudefontaine, France). The electrodes of both clinical centers had between 8 and 18 contacts per electrode, a diameter of 0.8 mm, 2 mm contact length and separated from each other by 1.5 mm. We implanted between 89 and 223 SEEG contacts per patient (total contacts recorded: 1590; mean of 159 contacts per patient, SD ± 55). To determine the exact location of each electrode and contact, a CT-scan/MRI data fusion was performed for each patient. SEEG signal from Marseille center was recorded on a digital system (Natus Medical Incorporated) with sampling at 1024 Hz or more with 16-bit resolution, a hardware high-pass filter (cutoff = 0.16 Hz), and an antialiasing low-pass filter (cutoff = 340 Hz). Data from Bucharest hospital was recorded with a Natus Quantum 128 system (Natus Neuro, Middleton, WI), sampling rate at 4096 Hz, 24-bit resolution and a hardware high-pass filter at 0.08 Hz.

### Independent component analysis

Because this work is focused on the analysis of hippocampal sources, we selected for ICA only the electrodes targeting this structure. ICA was run on the continuous traces of each electrode of each patient separately. We thus analyzed a total of 15 electrodes (N=15) obtained from the 10 patients.

ICA aims to solve the ‘cocktail party’ problem by separating N statistically independent sources that have been mixed in M recording sensors. It is a blind source separation methodology, as the spatial distribution and time-courses of the sources are unknown. To identify the sources, ICA assumes that they are immobile in space, i.e., that the proportional contribution of each source to every sensor is the same throughout the recording session. Each recorded signal *u*_*m*_(*t*) is thus modeled as the sum of *N* independent sources (*s*_*n*_(*t*)) multiplied by a constant factor (*V*_*mn*_):

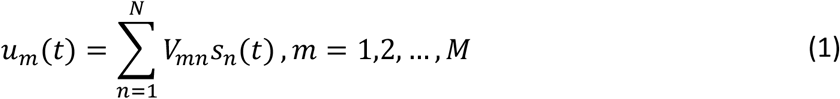

where *u*_*m*_(*t*) is the SEEG data, *V*_*mn*_ the ICA weights or spatial distribution of each source, *M* the number of sensors, *N* the number of sources and *s*_*n*_(*t*) the obtained independent components (“SEEG-ICs”).

In this work, we obtained as many components as sensors per electrode (N=M), without applying any dimension reduction (Artoni et al., 2018). We applied ICA on the continuous data, seeking one mixing matrix per electrode. We used FieldTrip (Oostenveld et al., 2011) to compute ICA based on the infomax algorithm, which aims to minimize the mutual information between components (Bell and Sejnowski, 1995), implemented in EEGLAB (Delorme and Makeig, 2004). Although only some of the SEEG-ICs were putative neuronal sources, we did not discard any component at this point.

### Analysis of event-related potentials

Only trials associated with the correct behavioral response were considered in the analysis. For each SEEG-IC, we checked if they had a significant ERP in both the “old” and the “new” conditions. To do so, we tested if each time point across trials was significantly different from zero with a t-test, obtaining a *t*- and *p* values for each time-point. Then, we corrected these tests for multiple comparisons using a local false discovery rate (LFDR; Benjamini and Heller, 2007) on the t-values with a threshold of 0.2 (Pizzo et al., 2019). LFDR assumes that the distribution of t-values is Gaussian, considering as significant those values that stand out from the distribution. To have a better estimation of the distribution, we grouped all the t-values across components of each electrode, obtaining a single threshold per electrode. To remove artifactual single points, we selected only those points during the first second after the stimulus and we imposed a minimum number of consecutive significant time samples (10 ms in this work).

To assess if the responses of the components differed in amplitude during “old” vs. “new” trials, we repeated a similar analysis across conditions. For each component and time-point we performed a t-test across trials between the amplitude of the ERP in old versus new conditions. Then, we corrected the t-values for multiple comparisons using LFDR on the t-values of each dataset (i.e., all the components of one electrode). In this way, we identified statistical differences at the single electrode level.

### Identification of hippocampal components

We focused on the putative hippocampal SEEG-ICs that were responding to the protocol. To identify these components from among all the obtained sources, we first selected the signals with a significant ERP. Then, we considered only the components with a local spatial distribution that was maximal in the hippocampus (i.e., those for which the ICA distribution presents a strong decay across sensors). After visual inspection, we identified two components that had the same ERP across electrodes. We labeled these components “Hc250-IC” and “Hc600-IC” based on the latencies of their responses. The former was present in 12 of 15 electrodes, while the latter was found in 13 electrodes. We only considered the time courses of Hc250-IC and Hc600-IC for further analysis. As ICA does not ensure the correct polarity and amplitude of the sources, we reversed the components when needed to match the same polarity across electrodes and z-scored the continuous traces of each component to facilitate the comparison across subjects.

### Spatial distribution across all recording sites

To test whether other brain regions were contributing to the SEEG-ICs components, we repeated the ICA now including all the recording sites in each patient. The computation of a single ICA for all the recordings may affect the resultant time-courses. Low variance hippocampal components may not be singled out, as the addition of many signals far from the hippocampus decreases the relative contribution of this source to the whole dataset. Moreover, the number of sensors and locations strongly differs across patients, hindering the inter-subject comparison. Therefore, we performed the analysis in an iterative manner, including only one additional contact at a time and evaluating its contribution to the SEEG-ICs. At each iteration, we computed a new ICA on the combination of the original electrode (i.e., the electrode targeting the hippocampus) and one extra contact. We then computed the zero-lag correlation between the original SEEG-IC signal (only the hippocampal electrode) and the new SEEG-ICs (combination of the hippocampal electrode and an additional contact), selecting the component with the highest correlation. This component would represent the same neuronal source in both datasets. Then, we estimated the relative contribution of the additional sensor to the component as the ratio between the ICA weight at the new location divided the highest ICA weight. A value close to 1 would imply that the additional contact strongly contributes to the SEEG-IC, suggesting that the neuronal source is nearby, while a value close to zero would indicate that the additional contact is relatively far from the source. This allows the analysis of the approximated spatial distribution of the components on the whole sampled brain, while minimizing the impact of including more contacts on the estimation of source time courses.

### Group analysis of event-related potentials

In order to test for a significative response at the group level, we performed a nonparametric permutation test corrected with cluster-based Statistics (Cohen, 2014). We computed the averaged ERP for each electrode and condition. Then, for each time point we computed a t-test against zero between the ERP values across electrodes in each condition. We kept the t-values of those points with a p-value lower than 0.05. These are the uncorrected values of significance. To correct for multiple comparisons, we selected clusters of significance, i.e., group of consecutive time points with a significant p-value. We assigned to each cluster the sum of the absolute t-values inside the cluster (absolute to also consider negative differences). We computed N=2000 surrogate datasets, by randomly shifting the ERPs of each electrode. The duration of each ERP was 2 seconds, including 500 ms of baseline, and the minimum shift of each surrogate was 250 ms. This way, the ERP signals remained the same, but the temporal alignment between ERPs was broken. We repeated the cluster procedure for each surrogate, keeping only the cluster with maximal t-value at each iteration. Any significance found in these surrogates would be by chance. Finally, we tested whether the t-values of our original clusters were significantly higher that the t-values of the surrogate analysis. The threshold of significance was set at the ninety-fifth percentile of the distribution of surrogate values (p-value < 0.05). The same procedure was followed to compare the amplitudes of the responses across conditions. In this case, the t-test was computed between conditions instead of against zero.

### Raster plots

We analyzed the trial-to-trial variability of the responses using raster plots. We selected all trials across electrodes and conditions and sorted them based on the RT to the stimulus. Then, we stacked them in a single matrix with dimensions time x number of trials. To assess whether the ERP was related with the patients’ response times, we correlated the latency of the ERP onset with the RT. First, we created super trials of 50 trials pooled across electrodes and conditions to improve the signal to noise ratio (Despouy et al., 2020; Hebart et al., 2018). We tested super trials of different sizes (in numbers of trials) without observing noteworthy differences in the main result. The onset latency of each super trial was estimated using the median rule, i.e., as the first time point with an amplitude higher than the median of the baseline plus 2.3 times the interquartile range (Letham and Raij, 2011). Finally, a Pearson correlation was applied between the onset latency and the averaged response time of each super trial.

### Detection of slope change points

To better characterize the dynamics the Hc600-IC response, we modeled the averaged ERP for each electrode and condition as a combination of multiple linear segments. The locations of the intersections between segments were automatically selected using the Matlab (Mathworks, Natick, MA) function *findchangepts*. This function identifies the points where the mean and the slope of the signal change most abruptly (Haynes et al., 2017). The total number of sections was given by a parametric threshold which imposes the minimum required improvement in the residual error for each change point. The residuals are strongly related with the signal to noise of the ERP, which was different for each electrode. Thus, we manually adjusted the threshold value between 0.5 and 3 for each ERP, until the response was clearly modeled with a minimum number of changepoints. The same value was used in both conditions. We determined the t2 value as the change point with maximum amplitude between 200 and 600 ms. Then, the t1 latency was selected as the first change point with a local minimum in amplitude before t2.

## RESULTS

### Segregation of SEEG time-courses into current generators

We performed electrophysiological recordings from the hippocampus in 10 patients during a recognition memory task (Figure 1a, see Methods). Each patient had between 10 and 20 electrodes in the whole brain, and we selected a total of 15 electrodes (N=15) implanted in this structure among all participants. The stereotactic electrodes had between 8 and 18 contacts that covered an entire path from the lateral to medial brain areas (Figure 1a). For each electrode, we applied the ICA source separation technique to segregate the recordings into the main sources contributing to the SEEG activity (Figure 1). Four different independent components (SEEG-ICs) were identified in most of the electrodes. The spatial profile of each component (Figure 1c) reflects its contribution to each SEEG contact, allowing to infer the location of the component’s source. The first component (labeled ‘Cort-IC’) is mainly visible on the lateral contacts of the electrode with a progressive reduction towards deeper structures, and presumably reflects a source located in the lateral temporal cortex. Two components had their maximal contribution inside the hippocampus and were labeled as ‘Hc250-IC’ and ‘Hc600-IC’ based on the latency of the ERP (Figure 1d). The former had its highest amplitude at ∼250 ms, while the latter, with a peak at ∼400 ms, may be related with the “hippocampal P600” commonly identified in visual recognition tasks (Barbeau et al., 2017; Trautner et al., 2004). Both components presented a narrow spatial profile with very low contributions from contacts outside the hippocampus, strongly suggesting that these components represented local sources inside this structure. The fourth component represented a remote source far from the electrode (‘Rem-IC’), with a similar contribution to all the contacts. The local components (i.e., Cort-IC, Hc250-IC and Hc600-IC) had a significant ERP compared to baseline (t-test across trials against zero, corrected with LFDR, see methods), but Rem-IC did not (Figure 1d).

**Figure 1:**
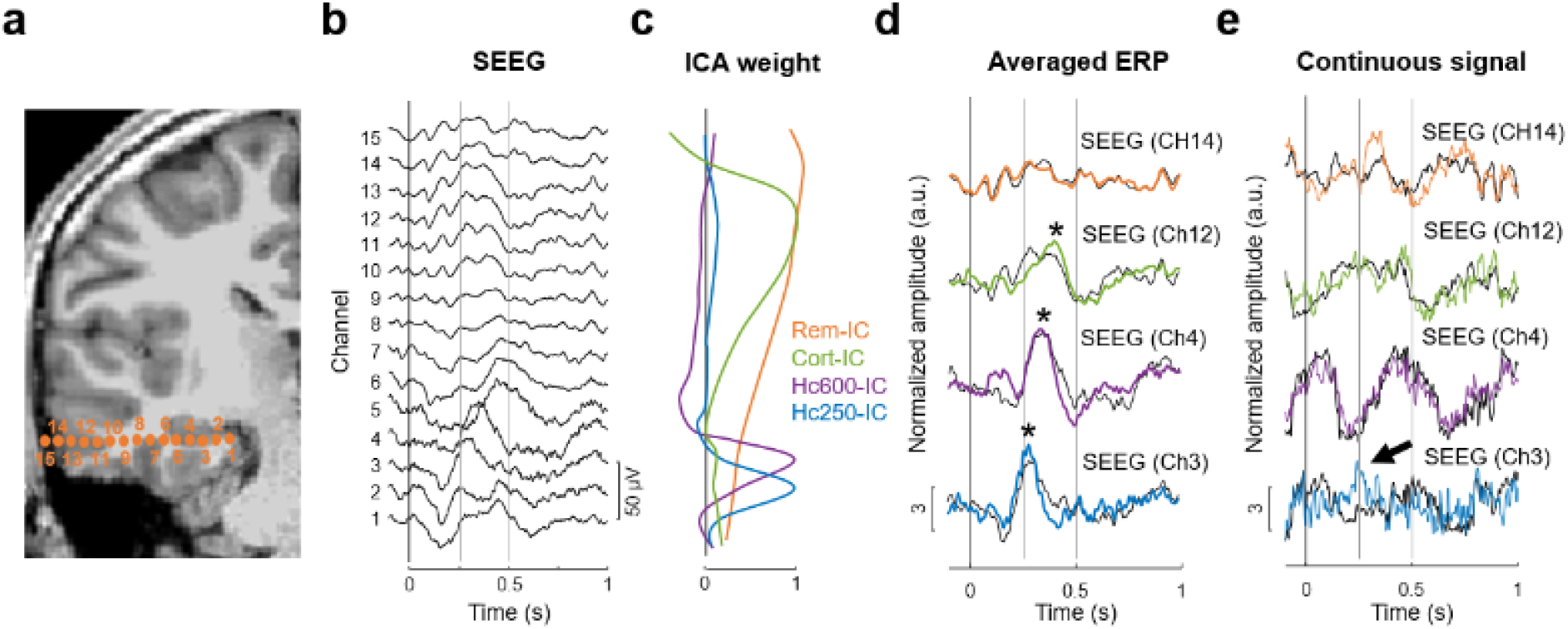
Separation of brain sources in SEEG with ICA in one patient. a) MRI (3D T1) with reconstruction of SEEG electrode for patient 1. The location of each recording site is represented with orange points. b) Averaged ERP for “old” (i.e., previously seen) items at each recording site (monopolar reference). c) Spatial profile of the SEEG-ICs across the electrode, representing the contribution of the SEEG-ICs to each sensor. d) Averaged ERP of each IC-EEG for “old” responses (color-coded traces; * p<0.05, t-test across trials against zero) superimposed with the SEEG channel (panel b) with maximal contribution from each SEEG-IC (black traces). e) Single trial response for SEEG-IC and SEEG. The arrow indicates a response to the stimulus that can be identified in SEEG-IC traces, but not on the original SEEG data.

The spatial profiles in Figure 1c show that the contribution of the different sources to the SEEG signals present an overlap, indicating that each sensor contains information from several components simultaneously. This is most noticeable in the hippocampus, where a single contact records activities from Rem-IC, Hc250-IC and Hc600-IC (Figure 1c, channel 3). ICA separates the time-courses associated with the sources, removing the contribution from other areas. For sources that are sufficiently separated in space, differences between SEEG-IC and SEEG may seem minor in the averaged ERPs (Figure 1d). However, these differences are remarkable in the continuous traces, where a response to the stimulus that is apparent in the SEEG-IC is hidden in the SEEG (Figure 1e, arrow). This contrast between SEEG-IC and SEEG is most prominent when the sources overlapped in space (Supplementary Figure 1). In this case, the activity of each source cannot be inferred from raw SEEG recordings (not even with bipolar montages, Supplementary Figure 1), and source separation methods are required to disentangle the time courses of the different components (Michelmann et al., 2018).

### Location and modulation of hippocampal sources

The number of local and remote sources retrieved with ICA varies across patients, as it strongly depends on the specific implantation scheme and the number of contacts per electrode. Thus, we focused our study on the two main hippocampal sources (Hc250-IC and Hc600-IC). Hc250-IC was present in 12 electrodes and its contribution to the dataset was small (explained variance across electrodes: 4,5%, SD ± 3,2%). Hc600-IC was identified in 13 of the 15 electrodes included in the analysis and represented an important contribution to the total variance of the data (mean explained variance: 29,2%, standard deviation: SD ± 14,2%). These components had a similar spatial distribution across patients; their maximal contribution was in contacts located inside the hippocampus, with little contribution from other sensors (Figure 2a). Within the hippocampus, the spatial profiles of both components presented were slightly different. Hc250-IC was recorded in deeper contacts than Hc600-IC in electrodes placed in the anterior hippocampus (Figure 2a, left). Intriguingly, this location was reversed in electrodes placed in the posterior hippocampus, where Hc250-IC appeared preferentially in more superficial sites (Figure 2a, right). This reversal may reflect rostral to caudal anatomical differences of the human hippocampus, well stratified in its anterior part and with complex folds of its substructures in more posterior sections (Andersen et al., 2006). Conversely, we did not observe notable differences neither in the spatial nor in the ERP between electrodes implanted in the left versus right hemispheres.

**Figure 2:**
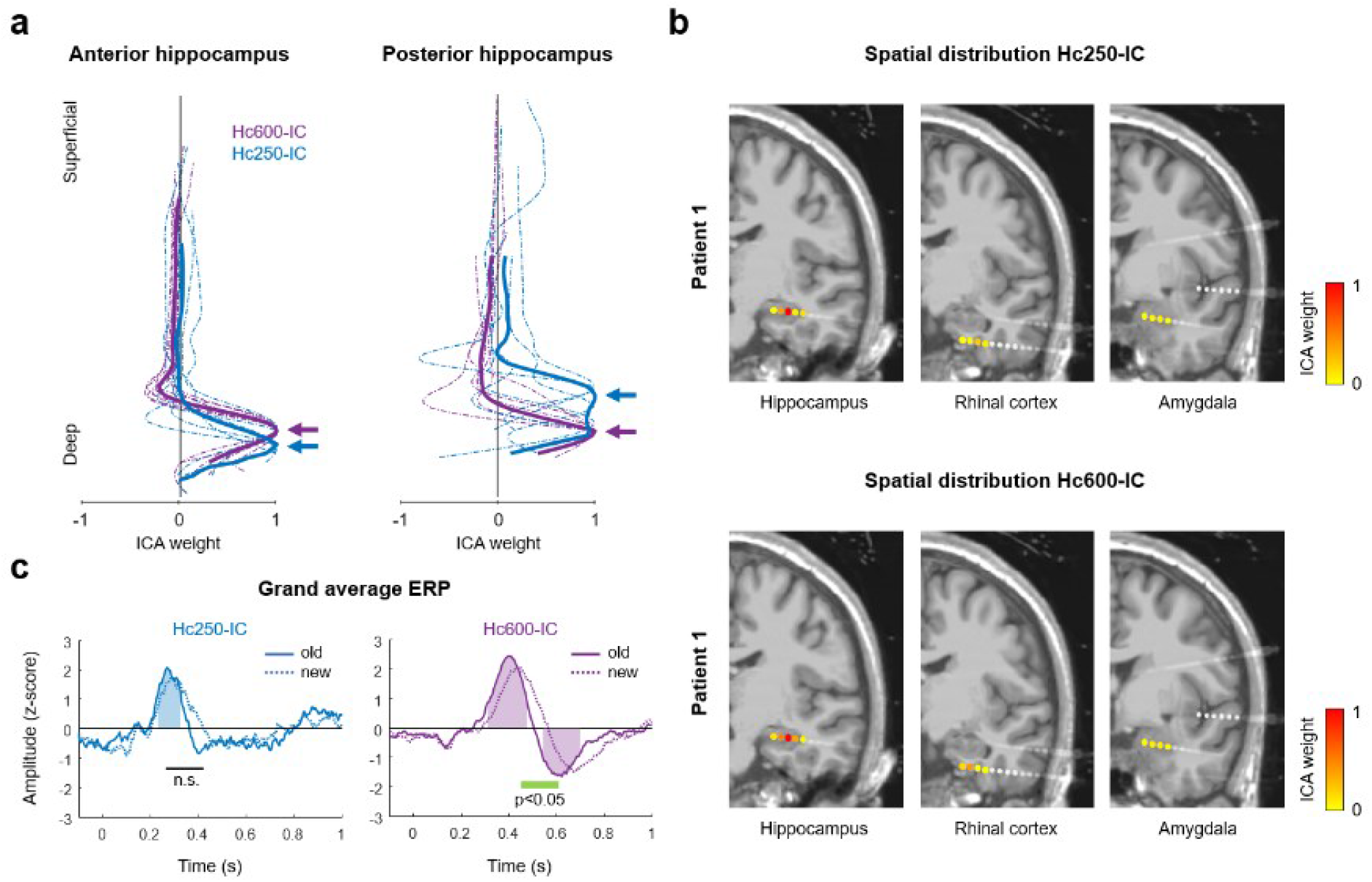
Two hippocampal sources during memory recognition. a) Spatial profile of the two hippocampal SEEG-ICs across electrodes located in anterior or posterior hippocampus. Dashed and solid traces represent the individual electrodes and their averaged value, respectively. Electrodes from different patients have been aligned based on the peak of Hc600-IC. Arrows indicate the location of the maximal value for the averaged profiles. b) MRI (3D T1) and reconstruction of SEEG electrodes for patient 1, with the spatial distribution of Hc250-IC (top) and Hc600-IC (bottom) across all the recording sites in one patient. The maximal contribution is in the hippocampus, with low values in all the other contacts, including those located in the amygdala and the rhinal cortex. c) Grand average ERP across electrodes contrasting “old” (solid traces) and “new” (dashed traces) items for Hc250-IC and Hc600-IC. Shaded areas illustrate the periods where the response to old trials is significantly different from zero, while the green line indicates the interval with differences between conditions (p<0.05, permutation test; n.s., not significant).

Although the spatial distributions of the SEEG-ICs presented a clear peak inside the hippocampus, this does not completely exclude the possibility that they reflect sources from a nearby region. To further test whether the sources were truly located in the hippocampus, we repeated the ICA now including all the contacts available in each patient (average n=159 contacts per patient). Computing a single ICA with all the sensors is not recommended because components with low variance (i.e., Hc250-IC) may be blurred among other sources. Thus, we performed this new analysis in an iterative manner, including only one additional contact at a time and evaluating its contribution to the SEEG-ICs. This allowed us to cover all the sampled brain areas while minimizing distortion of the time-courses identified in the restricted analysis (see Methods). For both Hc250-IC and Hc600-IC, the spatial distribution was in all cases maximal in the contacts within the hippocampus, with negligible contributions from other areas such as the rhinal cortex or the amygdala (Figure 2b). This result reinforces the interpretation of the hippocampus as the origin of the components.

The temporal profile of the components presented similar ERPs across electrodes (Figure 2c). There were no appreciable differences between components in the anterior and posterior hippocampus. Note that ICA does not ensure the correct polarity or amplitude of the sources. Thus, we reversed the components when needed and normalized them (z-score) to ease comparisons. The earliest response was in Hc250-IC, with a single peak at ∼260 ms (mean 257±42 ms across patients) post-stimulus during “old” items (Figure 2c, left) that was significantly different from zero in all electrodes (t-test across trials against zero, corrected with LFDR). The response to “new” items presented a similar timing as to recognized elements, but with significant differences in amplitude between both conditions in 6 out of 12 electrodes (old/new contrast; Supplementary Figure 2; t-test across trials between conditions, corrected with LFDR). However, there were no differences at the group level (p>0.05, permutation test). The response of Hc600-IC elicited by old items was characterized by two peaks of opposite polarity at 405 and 610 ms post stimulus onset (Figure 2c, right), similar to the standard hippocampal response in other memory tasks (Barbeau et al., 2017, 2008; Trautner et al., 2004). Interestingly, the visual comparison between “old” and “new” responses revealed that the changes were not in amplitude, but their temporal dynamics were different, with a slower activation time to “new” items. Therefore, we further explored the modulation of this component.

### Analysis of hippocampal dynamics

First, to confirm that the differences between conditions were induced by the memory paradigm, we tested whether the latency of the response was related to the behavioral response time (RT) to the protocol. We selected all the single trials for both “old” and “new” conditions and reordered them based on the RT, either for all the electrodes together (Figure 3a) or for each electrode separately (Figure 3b). No correlation between the raster plot and the RT could be visually appreciated. To quantify this observation, we grouped the data into super trials (i.e., the average of 50 trials with adjacent RTs pooled across electrodes and conditions) and estimated the evoked onset time of each one as the first time point with an amplitude significantly higher than baseline (see Methods). We then computed the correlation between such evoked onset time and the averaged RT of each super trial, with no significant relation between the timing of the Hc600-IC response and the RT (Figure 3c; correlation test, N=24 super trials, r^2^=0.006, p=0.71). This null result is in good agreement with previous studies, where the RT did modulate the response of the perirhinal cortex and the motor cortex, but had no discernable effect on the hippocampal response (Despouy et al., 2020).

**Figure 3:**
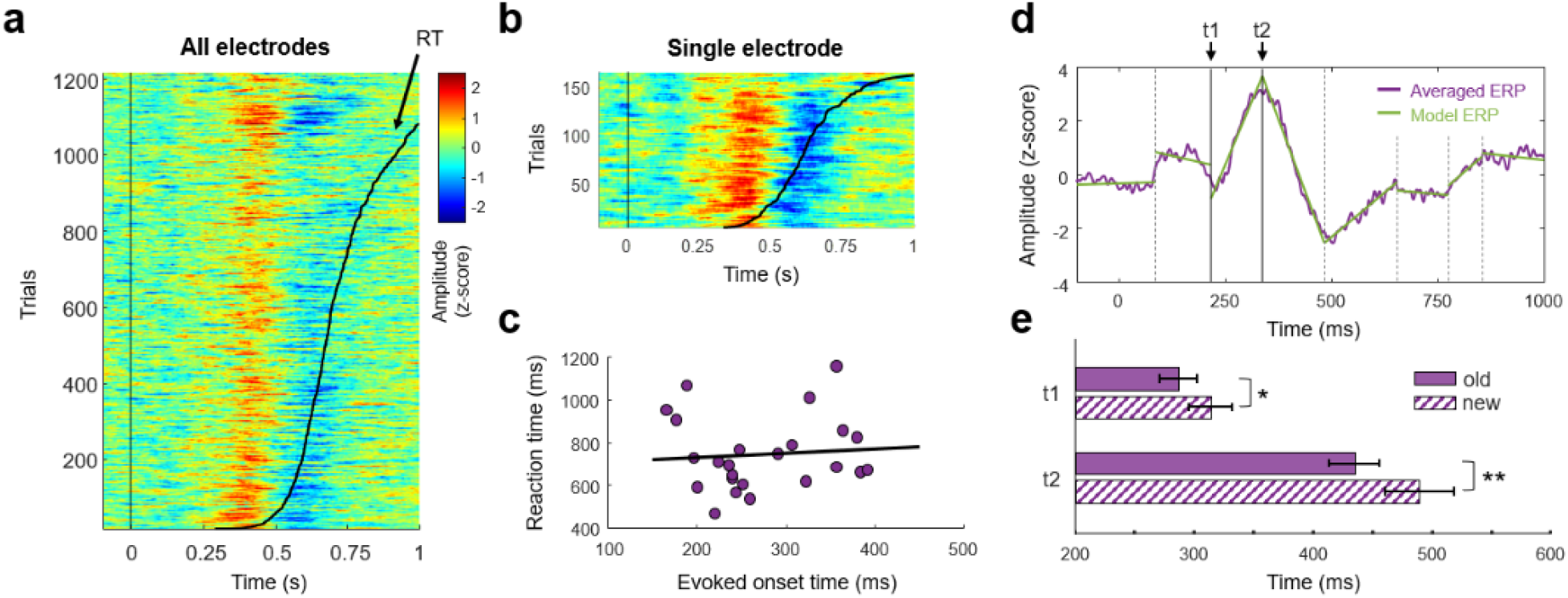
Comparison of Hc600-IC for old and new responses. a) Raster plot with single-trial ERP for old and new conditions across electrodes. Trials from all patients were ordered based on the response time (black curve). b) Raster plot for a representative participant. c) Correlation between the evoked onset time and the reaction time. Each dot represents data grouped across trials (super trial, see methods). d) Example of change points detection in the averaged ERP for old responses of one electrode. The ERP (purple trace) is modeled by several linear segments (green lines). The intersections between segments (vertical black lines) represent the time points with highest change in mean and slope. Two change points are selected: t1 where the amplitude starts rising and t2 where the amplitude is maximal. e) Comparison between t1 and t2 latencies across electrodes. (*/** p<0.05/0.01, paired t-test between conditions, N=13).

Second, to better analyze the properties of the temporal response, we modeled the averaged ERP of each condition and electrode as a combination of multiple linear sections (Figure 3d). The juncture between sections was statistically defined as the time-points where the mean and the slope of the signal changed most abruptly (see methods). We tested if the timing of these change points differed between conditions. Two points of the ERP were selected for each electrode: the latency at which amplitude starts to increase (t1) and the latency of the maximal response (t2). Responses to “old” images exhibited significantly earlier latencies at both t1 (mean latency ± standard error of the mean, s.e.m.: 287 ± 16.1 ms and 314 ± 17.9 ms for old and new responses, respectively; p<0.05, t-test across electrodes, N=13) and t2 (mean latency ± s.e.m.: 431 ± 21 ms and 490 ± 28.7 ms for old and new responses, respectively; p<0.01, paired t-test between conditions across electrodes, N=13; Figure 3e). This implies that the hippocampus processes memory recognition as soon as 290 ms post stimulus onset, around 120 ms before the latency of its maximal response.

## DISCUSSION

We described two different specific hippocampal sources recorded during a recognition memory task in humans. Patients with intracerebral electrodes implanted for focal drug-resistant epilepsy monitoring were asked to classify images as “old” or “new”. Using ICA on the SEEG recordings, we disentangled two hippocampal components (i.e., the spatial distribution maximum was inside the hippocampus) from other cortical and remote sources. The first source (Hc250-IC) had a low contribution to the total variance and presented an early response to the stimulus, with significant differences between memory conditions in 6 out of 12 electrodes. The second one (Hc600-IC) had a higher contribution to the total variance and was modulated by the memory condition in all cases, with faster responses for known images compared to “new” items. The earliest difference was at 290 ms, demonstrating that the hippocampus has an earlier involvement in memory recognition than previously reported.

### Identification of two hippocampal sources in SEEG

In this work, we have identified two different hippocampal neural generators, and, thanks to the use of ICA, separated their corresponding electrophysiological activity. The presence of several memory-related sources in the hippocampal formation has been previously reported (Barbeau et al., 2017; Ludowig et al., 2010). Ludowig et al., 2010, described two different generators, in the subiculum and in the posterior hippocampus, during an oddball task. These structures presented a similar ERP with the characteristic P300, but with different voltage gradients along the sensors, which indicated a local origin of the activities. It is unlikely that they are the components of our study, which we found in the anterior and posterior part of the hippocampus. Moreover, Hc250-IC and Hc600-IC had very different response time, in contrast with the generators identified by Ludowig and colleagues.

The two hippocampal components found in our study may be related to the different activities described in Barbeau et al., 2017. These authors observed that the evoked responses during memory and novelty detection had opposite polarities, raising the possibility of different sources for each response. It is possible that both sources were always present, but with one predominating over the other for each task. In this scenario, Hc600-IC would be the main generator modulated by memory recognition, while Hc250-IC could be related to a hippocampal source also involved in novelty detection. This would explain its faster activation (the response to novelty is believed to be faster than recognition: Barbeau et al., 2017; Norman and O’Reilly, 2003), and the absence of clear differences between conditions in memory recognition. The lack of differences between anterior and posterior hippocampus also supports this possibility (Poppenk et al., 2013). Further studies combining memory and novelty tasks will address this question.

Previous studies have suggested the presence of an early response (∼250 ms) from the hippocampus, modulated by stimuli repetition (James et al., 2009; Nahum et al., 2011; Raynal et al., 2020). This response would coincide in time with the Hc250-IC found in our study. However, they are unlikely to be generated by the same source. While in these studies the activity was related to memory encoding, we did not observe such changes for new elements. On the contrary, our results have the opposite tendency, with higher amplitudes during memory recognition at the single level. Moreover, the variance of Hc250-IC was quite low in the intracerebral recordings (4,5%), making its fingerprint on the surface likely to be negligible on raw EEG recordings (James et al., 2009; Raynal et al., 2020).

### Anatomical considerations of hippocampal current generators

The activity recorded at each sensor with SEEG reflects the summation of several close and distant sources (Buzsáki et al., 2012). The main features that determine the field potential of one region are the geometry and the degree of synchronization of the current sources (Herreras, 2016). Blind source separation methods such as ICA has been proposed as the solution to recover the time-courses associated to specific current generators (Herreras et al., 2022, 2015). Due to its versatility, ICA has been used to remove the reference signal in intracerebral EEG (Hu et al., 2007) and to separate neural sources in EEG (Onton et al., 2006; Tang et al., 2002), magnetoencephalography (MEG; Barborica et al., 2021; Malinowska et al., 2014) and local field potentials in rats (Fernández-Ruiz et al., 2021; Makarov et al., 2010; Torres et al., 2019). In this work, we innovatively used ICA to disentangle multiple generators in SEEG (Michelmann et al., 2018).

Several hippocampal generators have been described in animal studies (Benito et al., 2014; Fernández-Ruiz et al., 2021; Korovaichuk et al., 2010; López-Madrona et al., 2020), reflecting the inputs to different layers of CA1 and the dentate gyrus (DG). These two structures present a suitable anatomy to generate electric fields. In CA1, the most dominant generators are located in the *stratum radiatum*, with the input from CA3 through the Schaffer collateral (Benito et al., 2014; Korovaichuk et al., 2010), and in the *stratum lacunosum-moleculare* (Benito et al., 2014), where are located the synaptic outputs of the layer III of the EC. In the DG, the highest potential is generated by the projections from the layer II of the EC to the granular cells (Benito et al., 2014; Korovaichuk et al., 2010; Makarova et al., 2011). We speculate that the current generators in these two structures might be the origin of our components, with Hc250-IC related with the input of EC to the DG and Hc600-IC reflecting the computations in CA1.

The anatomy of the hippocampus differs from rostral to caudal (Andersen et al., 2006). The anterior section presents a relative clear distribution of the different layers, with the subiculum, CA1 and CA3 in the lateral part, surrounding the DG in between (Amaral, 1999). This results in well localized components, with similar topographies across electrodes (Figure 2a, anterior hippocampus). In good agreement with our hypothesis, the spatial profiles of Hc600-IC are deeper than those of Hc250-IC, which may relate Hc250-IC with lateral areas (but see below). On the contrary, the posterior section of the hippocampus has a different anatomy, with several folds of the granular layer, forming small “dents”. This geometry impacts the spatial distribution of its current sources, with summation and cancelation of currents caused by the opposite orientation of the cells through the gyrus (Buzsáki et al., 2012). Our results reflect the complexity of this area, with a huge variety in shape and location of the topographies of the two identified hippocampal sources across electrodes/patients (Figure 2a, posterior hippocampus).

It is important to remark the complexity of the hippocampal circuit to avoid simplistic interpretations. To exemplified this, the DG includes a dense inhibitory network, back projections from CA3 and several inputs from lateral and medial EC with different information (Andersen et al., 2006). Therefore, the identified generators cannot be linked to a single pathway or process. Moreover, the location of the maximal potential may not coincide with the origin of the current source (Herreras, 2016; Herreras et al., 2022). Due to the curvature of CA1, a synchronized synaptic input to the whole coronal layer (i.e., from the boundary with CA3 to the subiculum), would generate field potentials across the layer, whose sum would be maximal in the center of the curve. In this situation, an activation of CA1 could be detected inside the DG. This effect has been described in the DG, where the synaptic input of the EC was in the molecular layer, but its field potential was dominant in the hilar region (Fernández-Ruiz et al., 2013). Further information of the gradients of the field potentials across the hippocampus may help to identify the true origin of the components, for example, using microelectrodes to improve the resolution across SEEG sensors (Ulbert et al., 2001). Ultimately, a realistic computational model of the human hippocampus is necessary to understand the origin of the multiple source generators (Herreras et al., 2015).

### Early memory recognition system of the hippocampus

It has been suggested the existence of two different memory recognition systems in the brain (Despouy et al., 2020): one fast, linked to familiarity processes; and another slower, more related to memory recollection (Yonelinas, 2002). According to this theory, the fast system would respond between 200 and 300 ms after stimulus onset and it would involve a number of areas in the frontal, temporal and parietal lobes, leaded by the perirhinal cortex (Despouy et al., 2020; Gonzalez et al., 2015). This system may reflect our interaction with the environment, allowing us to rapidly react to any stimulus. The second system would be triggered at ∼450 ms after the stimulus, when the hippocampus and other areas of the medial temporal lobe are activated (Despouy et al., 2020). At this stage, more elaborated memories are retrieved with additional information.

As most of previous studies have focused their analysis on the main hippocampal response, the so-called hippocampal P600 (Barbeau et al., 2017, 2008; Despouy et al., 2020; Dietl et al., 2005; Trautner et al., 2004), it has been suggested that this structure cannot be involved in fast memory processing, which would occur much earlier. However, thanks to the use of ICA in SEEG recordings, we have found that the hippocampal dynamics during memory recognition are very complex, with at least two different generators contributing to the response. Our results, with the peak of Hc250-IC ERP at 250 ms and Hc600-IC presenting the earliest differential activity at 290 ms, challenges the vision of the hippocampus as a “late” structure.

We propose that both components (Hc250-IC and Hc600-IC) represent two steps of processing. The earliest component, Hc250-IC, presents similar delays as the N240 component evoked in the EC and in other mesial structures (Barbeau et al., 2017). Thus, it may reflect the input from the EC, the main entrance pathway to the hippocampus (Fernández-Ruiz et al., 2021; López-Madrona and Canals, 2021). This activity would already contain a pre-identification of known elements, realized in the perirhinal cortex (Despouy et al., 2020). The preprocessed information from EC may trigger the internal hippocampal circuit (linked to Hc600-IC), facilitating memory retrieval of those elements already recognized. This neuronal facilitation would be reflected in the ERP, with earlier latencies for old items (Figure 3e). Interestingly, we identified a similar facilitation pattern in magnetoencephalography (MEG) during the same task (López-Madrona et al., 2022). In that work, the combined activity of the hippocampus and the rhinal cortex presented earlier latencies for old items. Importantly, the origin of the MEG activity was validated using simultaneous MEG-SEEG recordings. Although the Hc250-IC responses suggest that the hippocampus process could be even faster in identifying known items, the absence of clear differences at the group level cannot confirm a direct link between the hippocampus and the fast memory recognition system. Further work is required, for example pushing the participants to their quickest answer and exploring the hippocampus together with other areas involved in the task. Overall, our results show that the hippocampus is not a late responding structure, but it is activated relatively fast during memory recognition (Despouy et al., 2020).

### Concluding remarks

The brain circuit of memory encoding and retrieval is an open question in neuroscience (Ferguson et al., 2019; Thompson and Kim, 1996). Our results provide new insights in the role of the hippocampus in memory recognition, linking the hippocampal activity to early stages of memory recognition. However, it remains unknown which is the specific contribution of the hippocampus to these fast processes. Future investigations will explore the role of the identified hippocampal sources in different memory processes, as well as the specific substructures contributing to the recorded activity. In addition, we have proved the efficacy of ICA to disentangle neuronal generators in SEEG. This opens new possibilities, not only in the analysis of the human hippocampus, but for all intracerebral studies. The use of ICA in other brain structures may reveal the dynamics of different but spatially overlapped structures, overcoming the limitations of traditional montages (Herreras et al., 2022, 2015). Importantly, it can be used in clinical applications with, for example, the potential to identify and separate the generators involved in epileptic networks (Barborica et al., 2021; Malinowska et al., 2014). Further research is granted.

## ACKNOWLEDGEMENTS

We would like to thank Cornel Tudor and Aurelia Dabu for performing the surgical procedures, Flavius Bratu, Camelia Lentoiu and Felicia Mihai for assisting with the patient investigation and data collection, Cristian Donos for the development of the computational framework for localizing the SEEG electrodes in common patient space. This study was performed thanks to a FLAG ERA/HBP grant from Agence Nationale de la recherche, ANR-17-HBPR-0005 SCALES and UEFISCDI COFUND-FLAGERA II-SCALES. This work has received support from the French government under the Programme « Investissements d’Avenir », Initiative d’Excellence d’Aix-Marseille Université via A*Midex funding (AMX-19-IET-004), and ANR (ANR-17-EURE-0029). The data were acquired on a platform member of France Life Imaging network (grant ANR-11-INBS-0006), supported in part by grants ANR-16-CONV-0002 (ILCB) and the Excellence Initiative of Aix-Marseille University (A*MIDEX).

## SUPPLEMENTARY MATERIAL

**Supplementary Figure 1:**
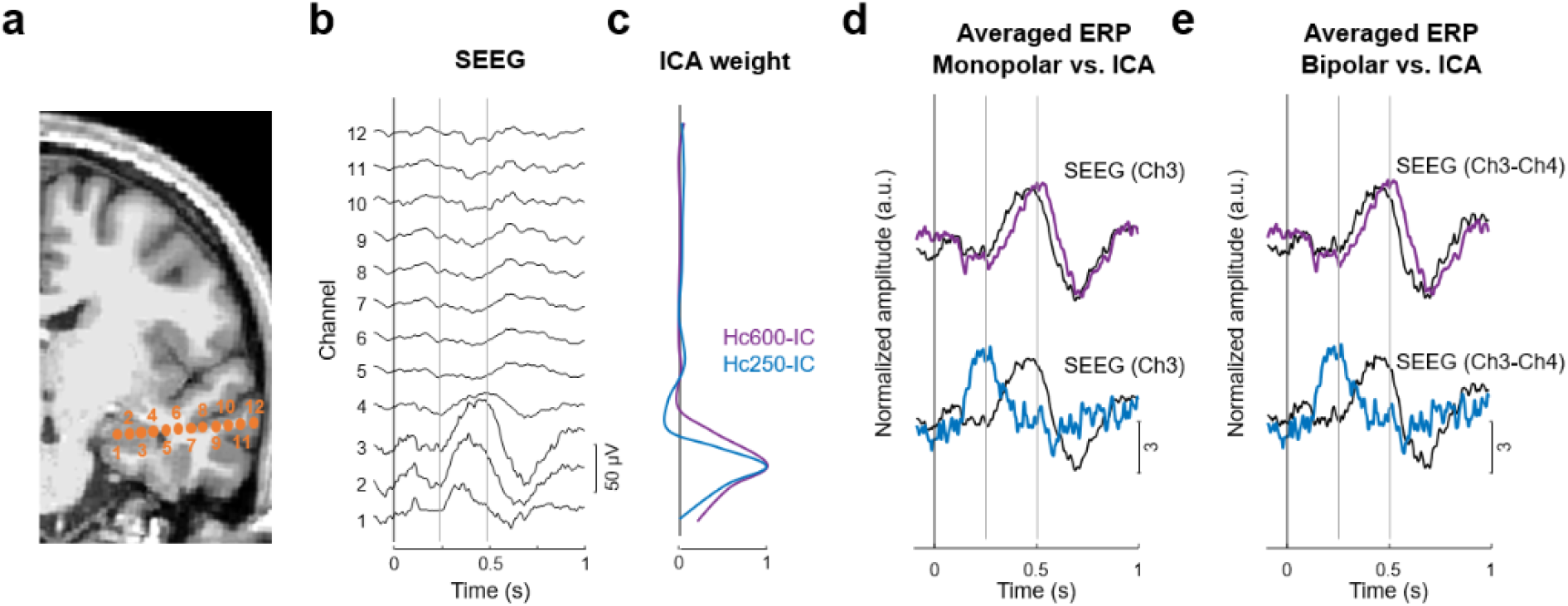
Separation of colocalized hippocampal sources in SEEG with ICA. a) MRI (3D T1) with reconstruction of SEEG electrode for one patient. The location of each recording site is represented with orange points. b) Averaged ERP for old responses at each recording site (monopolar reference). c) Spatial profile of the SEEG-ICs across the electrode. Both components are maximal at the same sensor, but their spatial profiles differ. d) Averaged ERP for old responses of SEEG-ICs (color-coded traces) superimposed with the monopolar SEEG at the location of maximal contribution from each SEEG-IC (channel 3, black traces. While Hc600-IC correlates with the SEEG response, the early response from Hc250-IC cannot be appreciated in the raw SEEG. e) Same as panel d, but with a bipolar montage for SEEG. As both sources are colocalized close to the channel 3, the local currents obtained with the bipolar montage represent the main current generator (Hc600-IC), which hides the activity from Hc250-IC. Note that ICA was always computed on the monopolar montage.

**Supplementary Figure 2:**
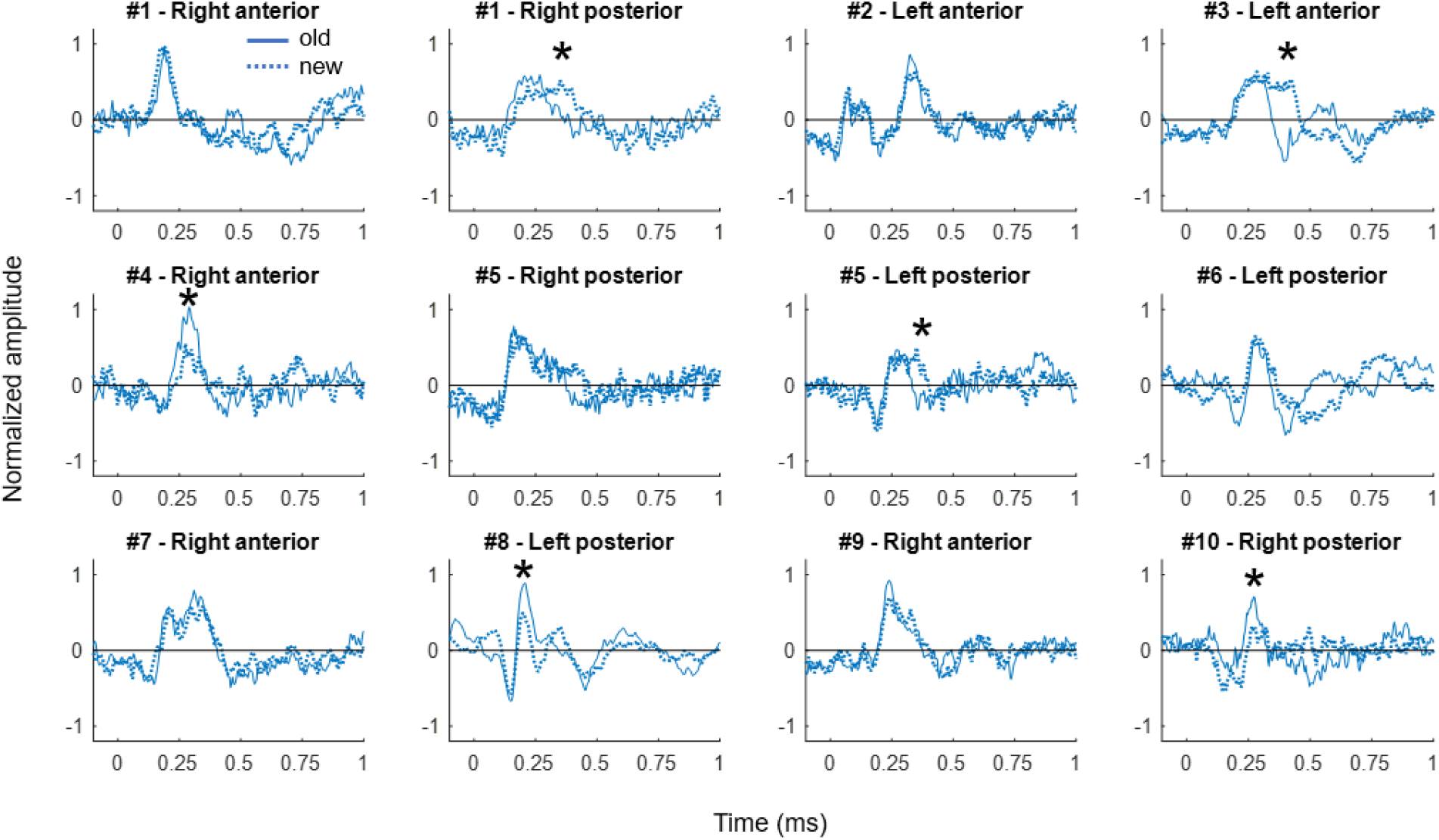
Single-case comparison between old and new responses of Hc250-IC. Each plot represents the averaged ERP of Hc250-IC during old (solid traces) and new (dashed traces) trials for each electrode. Hashes represent patient number and stars indicate significant differences in amplitude between conditions (* p<0.05, t-test across trials corrected with LFDR). Only 6 out of 12 electrodes presented a modulation to the memory protocol. In three cases, this difference was due to higher amplitudes after the presentation of old images at early latencies (electrodes 5, 10 and 12). In the other three electrodes, the responses were different, with the responses to the new images standing high during a longer period (electrodes 2, 4 and 7)

## Notes

### Competing Interest Statement

The authors have declared no competing interest.

